# Mobile genetic element-encoded putative DNA primases composed of A-family polymerase - SSB pairs

**DOI:** 10.1101/2022.12.02.518924

**Authors:** Phoebe A. Rice

## Abstract

Mobile genetic elements can encode a wide variety of genes that support their own stability and mobility as well as genes that provide accessory functions to their hosts. Such genes can be adopted from host chromosomes and can be exchanged with other mobile elements. Due to their accessory nature, the evolutionary trajectories of these genes can differ from those of essential host genes. The mobilome therefore provides a rich source of genetic innovation. We previously described a new type of primase encoded by *S. aureus* SCC*mec* elements that is composed of an A-family polymerase catalytic domain in complex with a small second protein that confers single-stranded DNA binding. Here we use new structure prediction methods in conjunction with sequence database searches to show that related primases are widespread among putative mobile genetic elements in the Firmicutes. Structure predictions show that the second protein adopts an OB fold (common among single-stranded DNA binding (SSB) proteins) and these predictions were far more powerful than simple sequence comparisons in identifying its homologs. The protein-protein interaction surface varies among these polymerase – SSB complexes and appears to have arisen repeatedly by exploiting partial truncations of the polymerase’s N-terminal accessory domains.

## 2 Introduction

Despite sharing similar core chemistry, the priming and elongation of DNA synthesis are usually carried out by different enzymes, termed primases and polymerases. Delegating these activities to separate enzymes most likely facilitates regulation of replication and improves the accuracy of the products – considerations that are more important in the replication of chromosomes than in the replication of mobile genetic elements. Nearly all known primases can be divided into just two large, structurally unrelated families: the DnaG family, which has a toprim motif at its core, and the AEP family, which has an RRM (RNA recognition motif) at its core (Aravind et al., 1998; Bergsch et al., 2019; Iyer et al., 2005). Although most primer-dependent DNA polymerases also contain the evolutionarily ancient RRM motif, the AEP – family primases contain other distinguishing features (Koonin et al., 2020; Raia et al., 2019). We recently reported an unusual mobile genetic element-encoded primase, CcPol-MP, that does not belong to either of the two canonical primase families (Bebel et al., 2020). Instead, CcPol-MP is a complex of two proteins: CcPol, which contributes an A-family polymerase domain that is responsible for the catalytic activity, and MP, which confers the ability to bind single-stranded template DNA. By combining database searches with structure predictions, we now outline a broader family of CcPol-MP-like primases, which we propose to call the PolA-SSB primases, and discuss how they may have repeatedly evolved new protein-protein contacts.

We discovered CcPol-MP as part of a broader effort to understand the core genes of the SCC (staphylococcal cassette chromosome) family of mobile genetic elements (Firth et al., 2018; Mir-Sanchis et al., 2016). SCCs are highly variable and carry a variety of accessory genes as cargo, the most well-studied being the methicillin resistance gene that creates MRSA (methicillin-resistant *S. aureus*) strains. SCCs share a conserved integration site and a core genetic locus. Central to the core locus is an operon encoding one or two site-specific DNA recombinases from the “large serine” family that can insert and excise the element (Misiura et al., 2013). Immediately preceding the recombinase operon is one that, while variable, always encodes a helicase (Mir-Sanchis et al., 2016). Related helicases (generally called “rep”) are found in the better-understood staphylococcal pathogenicity islands (SaPI), which are known to replicate as part of their life cycle (Fillol-Salom et al., 2018)(Qiao et al., 2022). In the SaPIs, the helicase is always preceded by, and sometimes fused to, a primase. In place of this primase, many SCC elements encode two ORFs in the same operon as the helicase: the 1^st^ is annotated as an A-family polymerase, and the 2^nd^ contains no previously identifiable conserved domains. We termed these CcPol for cassette chromosome polymerase and MP for middle protein (Bebel et al., 2020). The Aravind group had previously included CcPol as a member of the “TV-Pol” group of A-family polymerases that are encoded by transposons and <u>v</u>iruses and are closely associated with helicases (Iyer et al., 2008). They proposed that TV-Pols might act as primases due to their genetic context. However, CcPol lacks the N-terminal domain associated with many TV-Pols. We found that CcPol and MP co-purified and that the complex was indeed able to synthesize new DNA strands in a template-dependent but primer-independent manner. This primase activity required CcPol’s catalytic site: in the absence of MP, CcPol could only extend primers rather than synthesize them *de novo*. MP also conferred ssDNA binding activity to the complex.

These findings raised the question of how DNA polymerases are normally prevented from initiating synthesis *de novo* – that is, in the absence of a primer. The chemistry of joining the initial two nucleotides together is the same as the elongation step of adding a nucleotide onto the end of an existing primer: a 3’ hydroxyl attacks the alpha phosphate of a nucleotide triphosphate, displacing pyrophosphate. Therefore, the key differences between primases and primer-dependent polymerases must lie in the substrates and in their ability to utilize them: priming requires a single-stranded template and two (d)NTPs rather than a double-stranded primer-template duplex and a single (d)NTP. The modeling presented here suggests strategies that CcPol-MP may use to overcome barriers to *de novo* initiation.

Here we apply the new structural modeling capabilities of AlphaFold to address how MP confers ssDNA binding activity, how the two proteins interact, how widespread CcPol-MP type primases are and how variable they can be (Evans et al., 2022; Jumper et al., 2021). These questions were difficult to address previously in part due to difficulties in modeling MP and in deciding whether or not other open reading frames encode MP-like proteins. MP has no identifiable conserved domains and our earlier protein modeling attempts could only predict that it would be rich in beta strands. However, AlphaFold predicts that it adopts an OB fold, similar to that found in many single-stranded DNA binding proteins (SSBs) (Dickey et al., 2013). Our results suggest that primases based on an A-family polymerase paired with an SSB are quite widespread in the mobilome of Firmicutes, generally associated with a helicase and site-specific recombinase(s).

The polymerase component of these new putative primases appears to have evolved from a DNA Pol I and to have undergone multiple truncation events that entailed evolving new interactions with the cognate SSB. DNA Pol I contains 3 overall domains: an N-terminal 5’ to 3’ exonuclease domain, a central 3’ to 5’ exonuclease domain, and a C-terminal DNA polymerase domain (Raia et al., 2019). The polymerase domain, which resembles a right hand in shape, can be further subdivided into thumb, fingers, and palm subdomains. The catalytic active site is found in the palm subdomain, which contains an RRM (RNA recognition motif) at its core. The putative Pol component of some of the protein pairs described here contains little more than the RRM core (which may be vestigial and inactive) while others contain the full polymerase domain including key catalytic residues. The latter also contain part or all of the 3’ to 5’ exonuclease domain, and some contain an additional N-terminal segment as well. For simplicity, in this manuscript all of the ORFs containing putative polymerase domains are referred to as “Pol”. When Pol-SSB interactions could be confidently predicted, all but one example involved the variable N-terminal region of the Pol protein. Evolution of new interactions may have been facilitated by sticky hydrophobic surfaces exposed on the Pol subunit after random truncations within its N-terminal domains.

## 3 Materials and Methods

Our goal was not to carry out a comprehensive survey, but rather to find a broad variety of CcPol-MP like protein pairs from different bacterial species. To do so we used a mixture of BLAST searches (of NCBI and UniProt databases) and webFlaGs (Saha et al., 2021), using as bait our previously characterized CcPol protein, the helicases and recombinases associated with it, and related proteins from SCC-like mobile genetic elements that we had previously noted (Bebel et al., 2020; Mir-Sanchis et al., 2016). A broad range of hits was found by using proteins associated with CcPol as bait, then focusing on the proteins encoded directly upstream of the helicase. CcPol itself was less useful as bait because its Pol domain is so closely related to that found in “housekeeping” bacterial DNA Pol Is, whereas MP was problematic as bait because its sequence is poorly conserved. Examples for further analysis were chosen to maximize variety in the length and sequence of the Pol proteins.

A total of 17 protein pairs, all closely associated, if not co-operonic, with a helicase, were chosen for further analysis. Accession numbers are listed in Table 1. Two of these pairs are from SCC elements (an SCC*mec* type V and an SCC*mer*-like element) found in tandem in the same *S. aureus* strain, both of which we have demonstrated primase activity for ((Bebel et al., 2020) and Rodriguez, Pigli and Rice unpublished data). We noted that the *Bacillus weidmannii* proteins in Table 1 are identical, except for the 1^st^ 4 residues, to proteins found in many (but not all) *B. cereus* strains, suggesting relatively recent horizontal transfer. An example from *B. cereus* is listed in Table 1 but not included in other analyses here. The *Nialla Nealsonii* sequence appeared to have a premature stop codon near the N-terminus: removal of one nucleotide from the stop codon of a short upstream ORF created a single fused reading frame with a sequence homologous to and the same length as that of the *C. difficile* and *P. pinistramenti* sequences. It is unclear if this is an evolutionarily recent mutation or a sequencing error. The “fixed” version of the sequence was used here.

**Table 1.** Summary of ORFs described in the text.

| Species / strain | SSB - Pol interaction | Accession #s |  |  | helicase type | recombinase type |
| --- | --- | --- | --- | --- | --- | --- |
|  |  | Pol | SSB | Helicase |  |  |
| **Full polymerase domains** |  |  |  |  |  |  |
| Clostridiodes difficile DSM102860 | C-ter strand to N-ter | WP_077708779.1 | WP_077708780.1 | WP_077708781.1 | COG3378 | 1 Large serine |
| Paenibacillus pinistramenti | C-ter strand to N-ter | WP_138494364.1 | WP_138494363.1 | WP_138494362.1 | COG3378 | 2 Large serine |
| Salibacterium qingdaonense strain CGMCC 1.10 | C-ter strand to N-ter | WP_177195569.1 | WP_090927634.1 | WP_090927635.1 | COG3378 | IS66 family transposase |
| Metalysinibacillus jejuensis strain N25 | C-ter strand to N-ter | WP_108307365.1 | WP_108307366.1 | WP_108307367.1 | COG3378 | Y recombinase |
| Turicibacter sanguinis isolate MGYG-HGUT-1 | C-ter helix to N-ter | WP_187115462.1 | WP_147633390.1 | WP_147633389.1 | COG3379 | 1 Large serine |
| Niallia nealsonii | none predicted | WP_235852004.1 | WP_101176622.1 | WP_101176623.1 | COG3378 | 2 Large serine |
| Clostridium perfringens strain CHD32500R | beta hairpin to thumb | WP_208364035.1 | WP_075810340.1 | WP_208364034.1 | COG3378 | 1 large serine |
| Listeria newyorkensis strain FSL L7-1614 | C-ter helix to N-ter | WP_185389139.1 | WP_185389140.1 | WP_185389141.1 | COG3378 | 2 Large serine |
| Trichococcus pasteurii strain DSM 2381 | C-ter helix to N-ter | WP_086941901.1 | WP_086941902.1 | WP_086941903.1 | COG3378 | 2 Large serine |
| Bacillus cereus AND1407 (& others) | none predicted | WP_001192253.1 | WP_000989044.1 | WP_001293926.1 | COG3378 | 2 Large serine |
| Bacillus weidmannii strain FSL J3-0113 | none predicted | WP_064476008.1 | WP_064476009.1 | WP_064476010.1 | COG3378 | 2 Large serine |
| SCCmec type V ; S. aureus strain TSGH17 | barrel to N-ter | BAK57481.1 | BAK57482.1 | BAK57483.1 | COG3378 | 1 large serine (CcrC) |
| SCCmer; S. aureus strain TSGH17 | barrel to N-ter | BAK57462.1 | BAK57463.1 | BAK57464.1 | COG3378 | 1 large serine (CcrC) |
| **RRM domains** |  |  |  |  |  |  |
| Bacillus subtilis strain Bsp4 | none predicted | WP_101502618.1 | WP_101502617.1 | WP_101502616.1 | DUF927 | 1 large serine |
| Bacillus mycoides strain BPN51/1 | none predicted | WP_215600838.1 | WP_215600837.1 | WP_215600836.1 | DUF927 | 1 large serine |
| Paraliobacillus zengyii strain X-1125 | none predicted | WP_112182537.1 | WP_112182536.1 | WP_112182535.1 | COG3378 | 1 small & 1 large serine |
| Planococcus antarcticus DSM 14505 | none predicted | WP_065536350.1 | WP_065536351.1 | WP_065537334.1 | COG3378 | 1 large serine |
| Geobacillus vulcani PSS1 | no SSB | WP_051870794.1 | none | WP_031406462.1 | DUF926 | 1 large serine |

Individual protein and complex structures were predicted using the CoLab implementation of AlphaFold2 and Alphafold-Complex, with default parameters (Evans et al., 2022; Jumper et al., 2021; Mirdita et al., 2022). The ORF between the putative Pol and helicase ORFs was assigned as a putative SSB (MP homolog) if it was predicted to have an OB fold - this was true for all cases in Table 1 except *Geobacillus vulcani*, in which this ORF was absent. Most of the helicases associated with these PolA-SSB primases were annotated as containing COG3378 and/or the D5_N and Primase C_term conserved domains, as are the helicases associated with CcPol-MP pairs in many SCC elements and the “Rep” helicase from SaPI5 (Qiao et al., 2022). Despite frequent mis-annotation as “primase” it is important to note that these helicases do not contain primase domains – they are termed “primase C_term” because DnaG- and AEP-family primases are often found upstream of or N-terminally fused to helicases of this family. A few Pol-MP examples were associated with a DUF927-containing helicase, similar to the Cch helicase found in the some SCC elements (Mir-Sanchis et al., 2016) and the Rep protein of SaPIBov1 (Ubeda et al., 2007).

Pol-SSB complexes were modeled with 1:1 stoichiometry in agreement with our biochemical and preliminary cryoEM data for *S. aureus* SCC*mec* CcPol-MP. Because chromosomally-encoded bacterial SSBs are usually tetrameric, we also tested modeling of MP as a tetramer, but no interactions were predicted among the 4 copies.

The percent identity matrices in Figure S1 were calculated by clustal omega (Sievers et al., 2011). For the longer polymerase proteins, the calculations were done with the highly variable N-terminal domains removed (based on alignment of the predicted structures). The structure-informed sequence alignment of the larger polymerases shown in Figure S3 was made using Promals3d (Pei et al., 2008). Structure figures prepared using The PyMOL Molecular Graphics System, Version 2.0 Schrödinger, LLC.

## 4 Results

The ORFs examined were highly diverse in sequence (Figure S1). The predicted putative polymerase structures could be grouped into 2 overall categories: 5 with little more than an RRM motif and 12 with a full polymerase domain (Table 1; see Figure S2 for information regarding the estimated accuracy of each prediction). Pairwise sequence identity among the former varied from 10% to 54% and among the latter (for polymerase domains only) from 16% to 55%, except for the two polymerase domains from *S. aureus* which shared 79% identity. Only the *geobacillus vulcani* example lacked an SSB-containing ORF. Pairwise sequence identity among the SSB proteins ranged from 7% to 70%, with the majority of pairs sharing less than 20% identity. All of these examples are found in Firmicutes, although we did not purposefully limit our searches to that group of bacteria. However, their niches vary widely, from humans (e.g., methicillin resistant *Staphylococcus aureus and Clostridiodes difficile*) to pine litter (*Paenibacillus pinistramenti*), sea-salt pans (*Salibacterium qingdaonense*) and an antarctic lake (*Planococcus antarcticus*)

### 4.1 Overall comparisons of structural models

Five of the putative Pol proteins analyzed contained little more than a conserved central RRM motif with small, variable N- and C-terminal extensions (Figure 1). The RRM motif corresponds to the active-site containing palm subdomain of multiple polymerase families. In the A-family bacterial DNA Pol Is, strands 1 and 3 of this motif harbor two key aspartate residues that bind catalytic Mg^++^ ions. However, in the 5 structural models shown in Figure 1, only 2, those from Bacilli, retain even one of those key residues. These proteins are therefore unlikely to be catalytically active. RRM motifs are found in a broad variety of proteins and it may be that despite their similar genetic context to the other putative primases studied here (and the proven ones from *S. aureus*), these five ORFs perform a different function.

**Figure 1.**
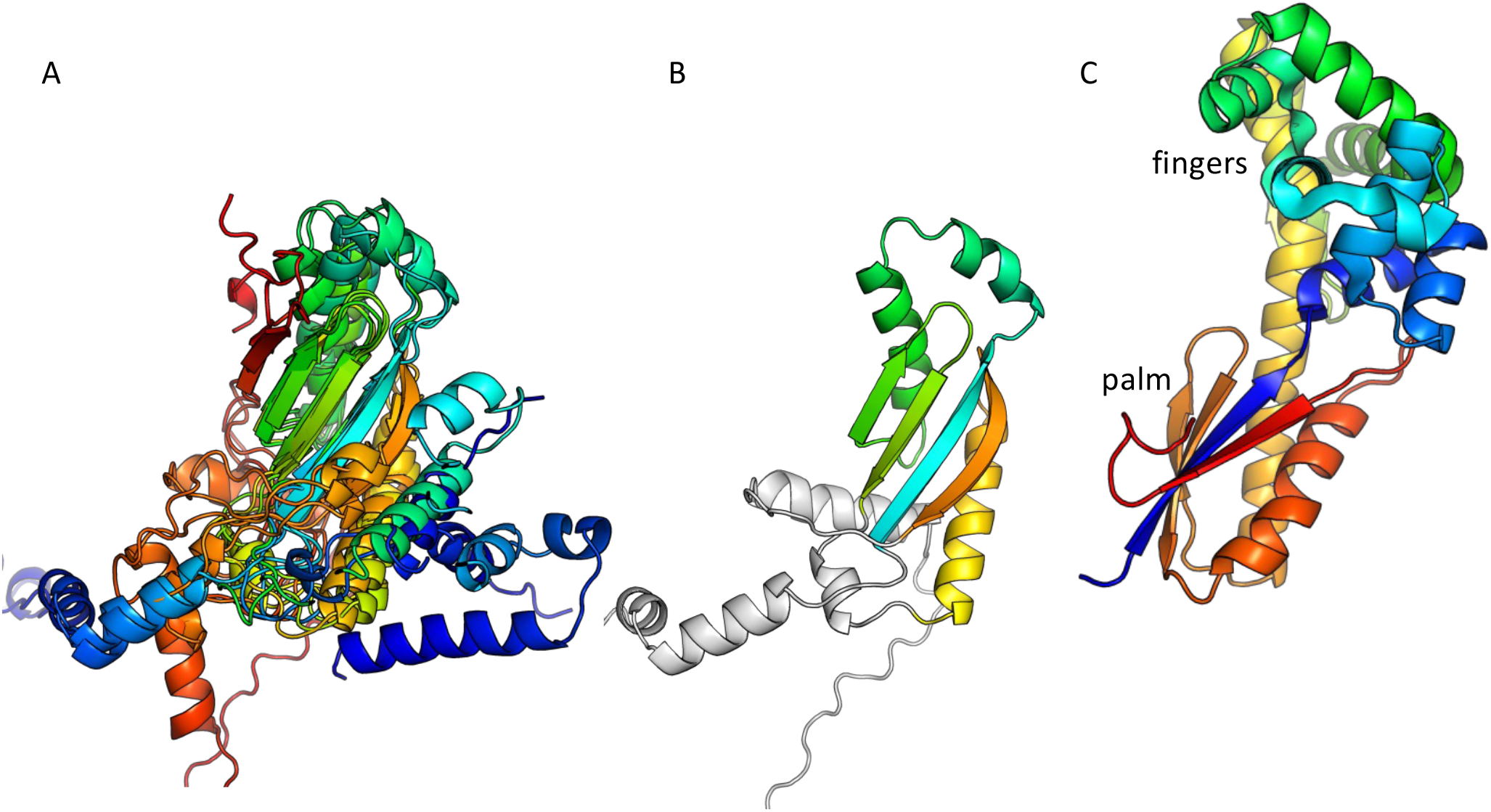
Predicted structures of the putative Pol proteins that contain little more than an RRM motif. A) All 5 examples listed in Table 1, colored blue to red from N to C, and superimposed according to the central beta sheet. B) The example from *B. subtilis* strain Bsp4 with with only the portion corresponding to an RRM motif colored. C) The palm and fingers subdomains of E. coli DNA Pol I are shown for comparison (1l5u.pdb; (Johnson et al., 2003)).

The remaining 12 putative Pol proteins were predicted to contain intact polymerase domains, including conserved Mg^++^ - binding active site residues in the palm subdomain and additional conserved residues in the functionally important helix O of the fingers subdomain (Figures 2 and S3) (Castro et al., 2009; Polesky et al., 1992). As shown in Figure 2, the predicted N-terminal regions varied. When a full 3’-5’ exonuclease was predicted to be present (7 models), key active site residues were also present (Figure S3)(Brautigam et al., 1999). Six of the examples with full polymerase domains were predicted to include a small additional N-terminal segment before the 3’-5’ exonuclease that may be a minimized relic of DNA Pol I’s large N-terminal 5’-3’ exonuclease domain, or may have been acquired through recombination. One predicted structure, that of the *C. perfringens* example, contains a full 3’-5’ exonuclease but no additional N-terminal segments. Five predicted structures, including the two *S. aureus* CcPols, include only part of the 3’-5’ exonuclease, yet confidence in the predicted fold of this partial domain was high except for the *Bacillus Weidmannii* example (Figure S2). The *S. aureus* CcPols also have a shortened thumb relative to the others and to DNA Pol I (Figure 2). *E. coli* DNA Pol I retains activity when its thumb is truncated, but shows reduced processivity and fidelity (Minnick et al., 1996).

**Figure 2.**
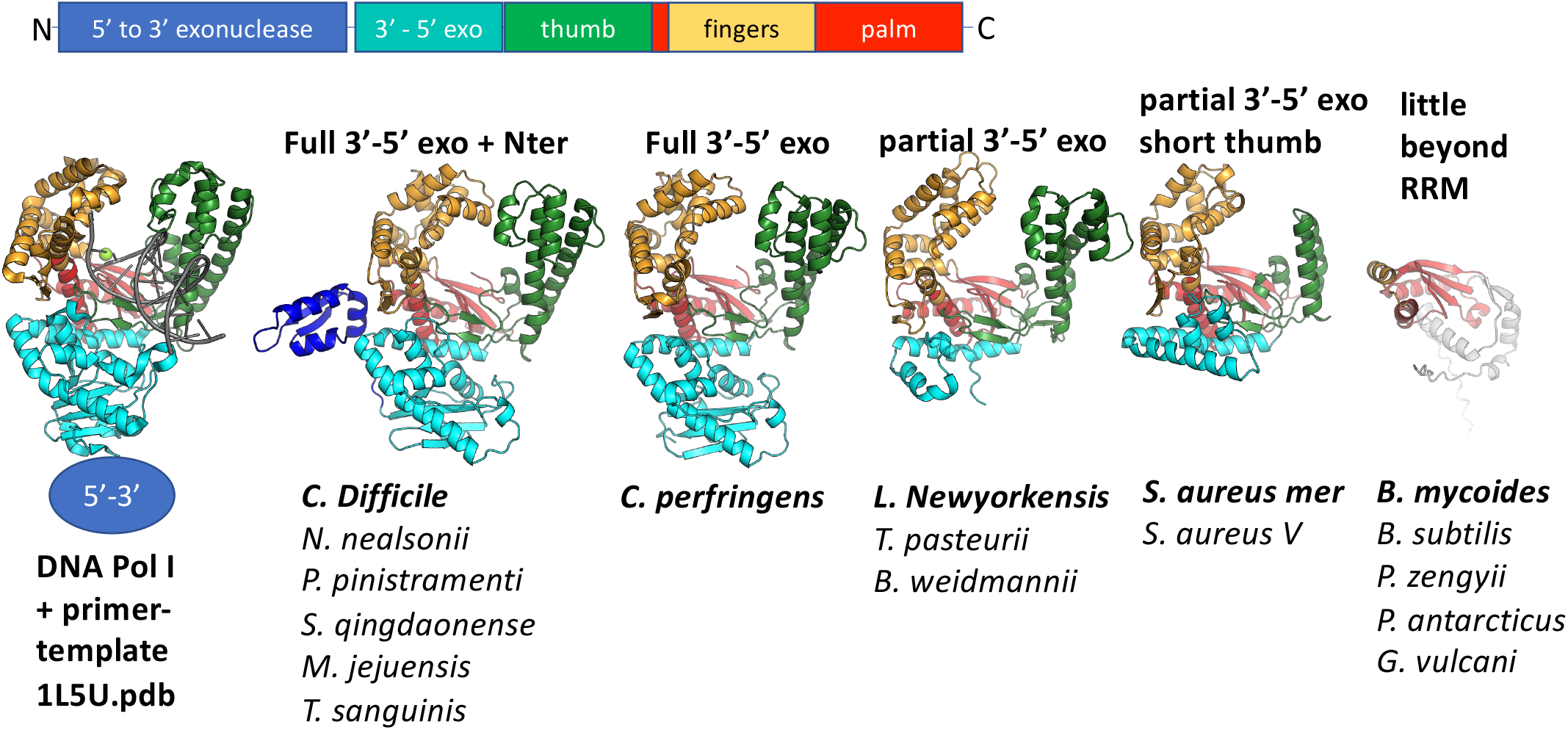
Comparison of predicted polymerase structures to bacterial DNA Pol I. Top: cartoon of the domains of DNA ol I. Bottom: representative structural models colored according to domain. In each panel, the bold label corresponds to the model shown, and those listed beneath it are very similar. A crystal structure of the Klenow fragment of Bacillus DNA Pol I with a bound primer-template duplex is shown for comparison (Johnson et al., 2003).

Although at the time of our initial studies of CcPol-MP (Bebel et al., 2020), we could not predict a structure for MP, AlphaFold now reliably predicts a small OB-fold beta barrel. This fold is commonly found in single-stranded DNA binding proteins (SSBs) (Dickey et al., 2013). For 16 out of 17 examples, the ORF sandwiched between the putative polymerase and the helicase was predicted to adopt this fold (Figure 3). In the *Geobacillus vulcani* case, we modeled the two ORFs directly upstream of the helicase, but the 1^st^ model was unrelated to our proteins and the 2^nd^ contained the RRM motif described above. Figure 3 also shows that the ssDNA-binding cleft of MP is predicted to have a positive electrostatic potential, as expected for a DNA binding protein. This structural model provides a good explanation for our previous observation that MP conferred ssDNA binding on the CcPol-MP complex (isolated MP was too poorly soluble for rigorous DNA binding assays). Based on these observations, we refer to this set of proteins as SSBs.

**Figure 3.**
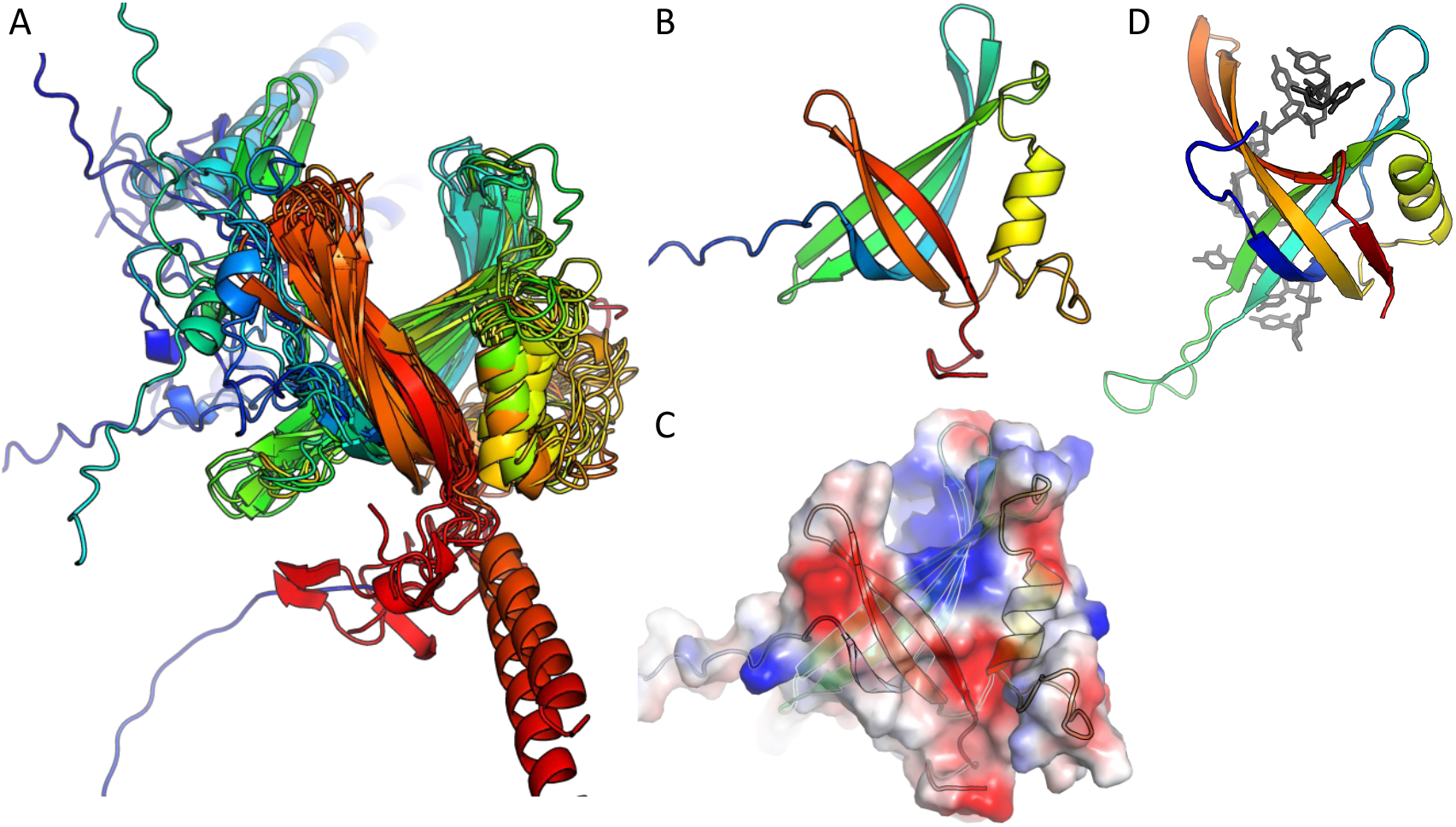
The predicted SSB subunits have an OB fold. A) Superposition of all 16 predicted structures, colored blue to red from N to C. B) The predicted structure of MP from *S. aureus* type V SCCmec. C) Semi-transparent surface of the model in (B) colored according to vacuum electrostatics (blue is positive; red is negative) (D) one subunit from E. coli SSB, with ssDNA bound, for comparison (PDB ID 1eyg; (Raghunathan et al., 2000))

### 4.2 Pol – SSB interactions

In agreement with our previous finding that *S. aureus* CcPol and MP (its cognate SSB) co-purify, AlphaFold predicted structurally plausible Pol – SSB interactions with low predicted alignment errors (implying high confidence) for most of the examples that included a full polymerase domain. The only exceptions were the *N. nealsonii* and the *B. weidmannii* pairs. For the latter, even the intramolecular predicted alignment errors were high for the N-terminal partial exonuclease domain (unlike the others with which it is grouped in Figure 2). Figure 4 shows the variety of predicted complexes.

**Figure 4.**
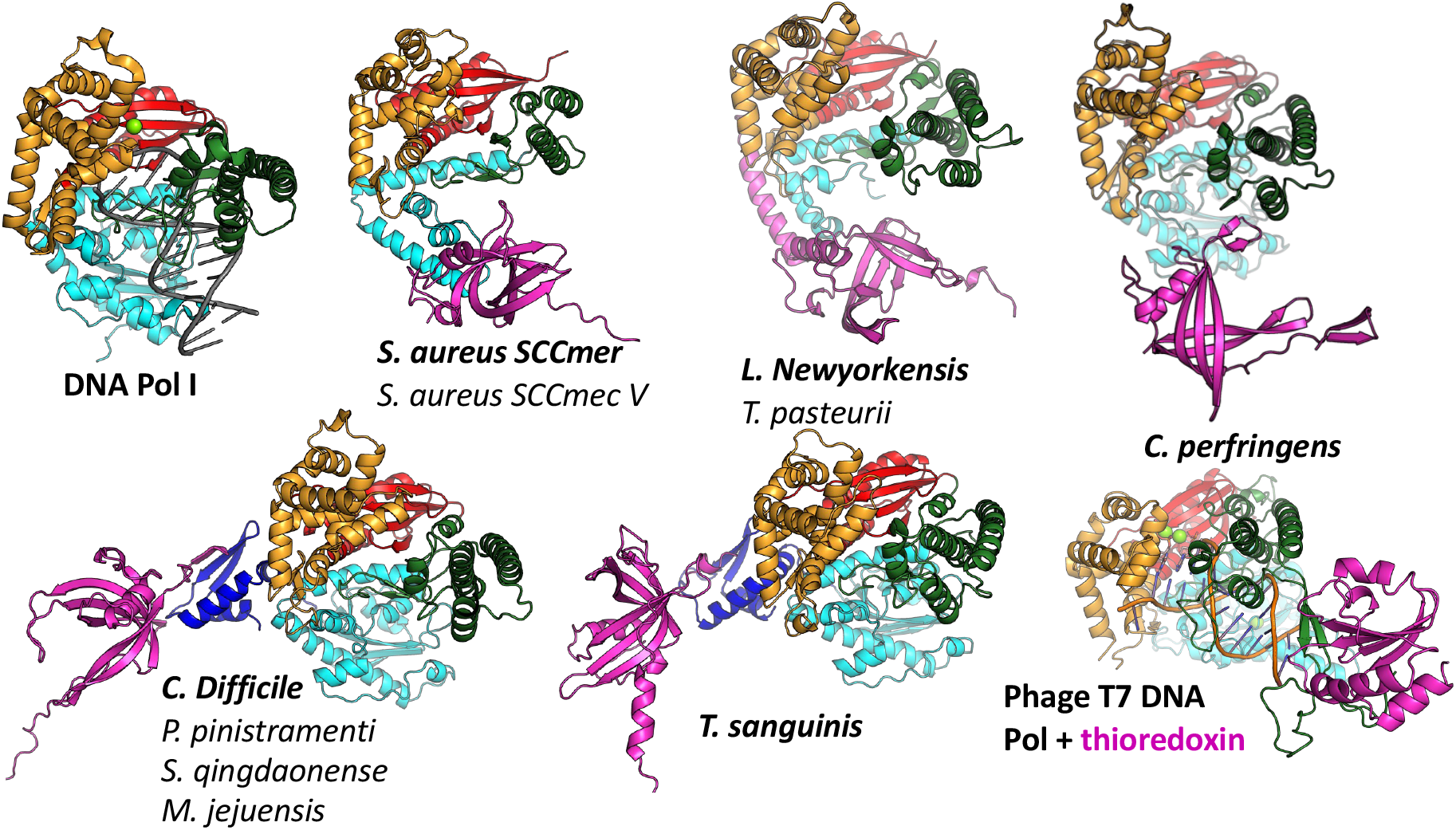
Predicted modes of Pol – SSB interactions. Models are colored as in Figure 2, with putative SSBs in pink. They were aligned according to their palm domains, except for T7 DNA polymerase, which was aligned to the C. perfringens model using the thumbs as guides. DNA Pol I (1l5u.pdb; (Johnson et al., 2003)) and T7 DNA polymerase (1t7p.pdb; (Doublié et al., 1998)) crystal structures are included for comparison. In each panel, the bold label corresponds to the model shown, and those listed beneath it are highly similar.

In most cases, the SSB is predicted to interact with the variable N-terminal regions (Figure 4) rather than with a more conserved segment of the polymerase. For those predicted to have only remnants of the 3’-5’ exonuclease domain, the SSB appears to have found two different solutions to the binding problem, both of which align the SSB’s positive cleft with the polymerase’s: in the *S. aureus* examples, the interaction is mediated by the barrel of the conserved OB fold, whereas in the *L. newyorkensis* and *T. pasteurii* examples, the interface is mediated primarily by a helical C-terminal extension found on these but not the other SSB models. For those 5 models that included a full exonuclease plus a small additional domain N-terminal domain, that N-terminal domain was predicted to mediate interactions with a C-terminal extension of the SSB protein: in four cases, the SSB adds an additional strand to the polymerase’s small beta sheet, and in the fifth case (*T. sanguinis*) a single-turn helix from the SSB docks into a similar location. Although these models appear to place the SSB rather awkwardly relative to the Pol domain, there is a flexible-looking linker between the very N-terminal domain and the 3’-5’ exonuclease that is likely to allow repositioning. The similarity of these latter five models, given that the polymerase proteins are no more than 55% identical to one another in sequence (Figure S1), enhances our confidence in them.

The *C. perfringens* example is predicted to use part of the Pol domain itself (the tip of the thumb) rather than an N-terminal region for SSB interactions (Figure 4). Although surprising, this predicted interaction is reminiscent of how phage T7 DNA polymerase (also an A-family polymerase) binds thioredoxin, which it exploits as a processivity factor (Bedford et al., 1997). The *C. perfringens* polymerase model is also the only one among the full-polymerase set that does not appear to have a partially truncated domain at its N-terminus: its N-terminus maps cleanly to the domain boundary between the two N-terminal exonuclease domains of DNA Pol I.

## 5 Discussion

The survey presented here shows that PolA-SSB type primases, which we previously discovered in certain *S. aureus* SCC*mec* mobile genetic elements, can be found a wide variety of Firmicutes. The PolA-SSB enzymes may represent a subset of the TV-Pol family previously suggested to function as primases due to their synteny with helicases, distinguished by their OB-fold-containing SSB subunit. Several lines of evidence suggest that all the PolA-SSB (and RRM-SSB) pairs described here are encoded by mobile genetic elements. First, the CcPol-MP pair that initiated our interest in this family are found on the SCC family of genomic islands of *S. aureus* (including many of the methicillin resistance-carrying SCC*mec* elements that create MRSA strains). Second, they appear to occur sporadically, rather than universally, in any given species. Third, at least in all of the examples listed in Table 1, one or more DNA recombinases are encoded just downstream of the helicase. In most cases, those recombinases belong to the “large serine” group of site-specific DNA recombinases, as do the CcrA/B/C recombinases of SCC elements (Misiura et al., 2013).

What might be the biological function of these primases? The simplest answer is DNA replication. However, the purpose of such replication is unknown, even for the SCC elements. One possibility is that the primase/helicase pair supports replication after excision from the host chromosome, which could enhance the efficiency of any mechanism of horizontal transfer to new hosts. Another possibility is that these proteins are responsible for synthesizing the 2^nd^ strand after horizontal transfer via natural competence or conjugation, both of which result in a single-stranded incoming donor DNA (Shen et al., 2022)(Blokesch, 2016). The latter possibility is supported by a recent report of horizontal transfer of SCC*mec* by natural competence (Maree et al., 2022). Stable incorporation of SCC*mec* into a new host chromosome required the presence of the recombinase genes. SCC-encoded replication machinery could promote the conversion of the incoming ssDNA to a duplex substrate for the recombinases. Alternatively, recent discoveries of numerous new systems that defend against invading DNAs (and are often encoded on mobile genetic elements) (Rocha and Bikard, 2022) raises the possibility that some or all of the PolA-SSB primases and their associated helicases are part of an uncharacterized type of defense system.

PolA-SSB primases presumably evolved from an ancestral DNA Pol I. What were the key changes that allowed a previously primer-dependent polymerase to initiate DNA synthesis *de novo*? The initial step in priming uses slightly different substrates than the elongation reaction: priming requires that the enzyme bind a single stranded template and two (d)NTPs, whereas for elongation, it must bind a primer-template duplex and one (d)NTP. The most relevant comparison between enzymes that can initiate *de novo* and ones that cannot is between the RNA polymerases (RNAPs) of bacteriophages such as T7 and the chromosomally-encoded bacterial DNA Pol I enzymes, all of which belong to the A family of polymerases (Dai and Rothman-Denes, 1998; Kennedy et al., 2007). Our modeling suggests that CcPol-MP and these RNAPs use analogous strategies to overcome barriers to initiating DNA synthesis *de novo*. Figure 5 and the sequence alignments in Figure S3 show that in T7 RNAP, CcPol and the other polymerase domains modeled here, the last helix of the thumb subdomain is one turn shorter than in DNA Pol I. The additional turn seen in DNA Pol Is is likely to sterically interfere with the triphosphate group of an initiating nucleotide, but not with the backbone of an elongating primer-template duplex. T7 RNAP also features an additional N-terminal domain not found in DNA Pol Is that recognizes cognate transcriptional promoters and orients the single stranded template in the polymerase active site (Kennedy et al., 2007). The SSB component of our PolA-SSB primases could similarly bind and orient the single stranded template, and in fact, we have already demonstrated that it confers ssDNA binding activity in the CcPol-MP case.

**Figure 5.**
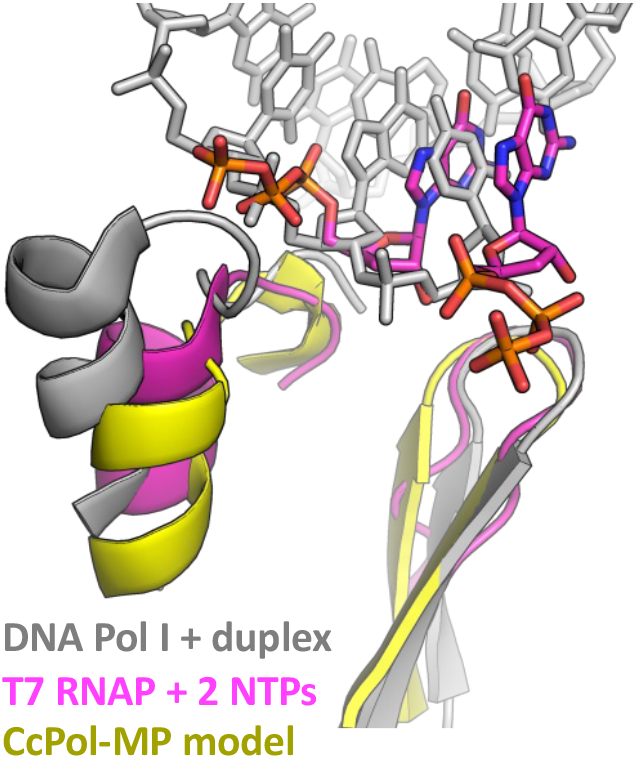
DNA Pol I may block the initial triphosphate. The last helix in the thumb and the central beta hairpin of the palm subdomains are shown. This superposition suggests that an initial triphosphate might make steric clashes with the last of the thumb of DNA Pol I’s helix (and/or the turn leading into that helix). That helix is shorter in T7 RNA polymerase and in all the full-polymerase models reported here. The Pol I coordinates were taken from PDBid 1l5u (Johnson et al., 2003) and the T7 RNAP coordinates from 2pi4 (Kennedy et al., 2007).

How did the PolA-SSB interaction evolve and why is it so variable? The N-terminus of the polymerase, which is found at the beginning of the 3-protein operon, appears to have been randomly truncated at different positions. Truncations in the middle of a folded domain would have left an exposed hydrophobic surface that the SSB may have been able to interact with: presumably weakly at 1^st^, then optimized through evolutionary selection. Only the *C. perfringens* example does not follow this trend: its N-terminus maps to a domain boundary, and its SSB is instead predicted to bind the tip of the (intact) thumb subdomain. We also noted that most of the predicted interactions are mediated by the variable C-terminal tail of the SSB component, and that the chromosomally-encoded canonical bacterial SSB protein also uses a different C-terminal extension to bind other replication-related proteins (Shereda et al., 2008). Further study with a far larger sequence data set would be needed to determine if the individual PolA-SSB pairings described here arose through single, independent truncations of the Pol protein, or if they have arisen sequentially through progressive random truncations of the Pol protein followed by re-optimization of new Pol-SSB contacts. The evolutionary paths of proteins that are only required for the horizontal transfer and/or maintenance of mobile genetic elements may be interestingly different from the evolutionary paths taken by essential proteins which cannot “tunnel” through non-functional intermediate sequences.

## Acknowledgements

This work was funded in part by NIH / NIGMS R01 GM121655.

---

**Figure S1.**
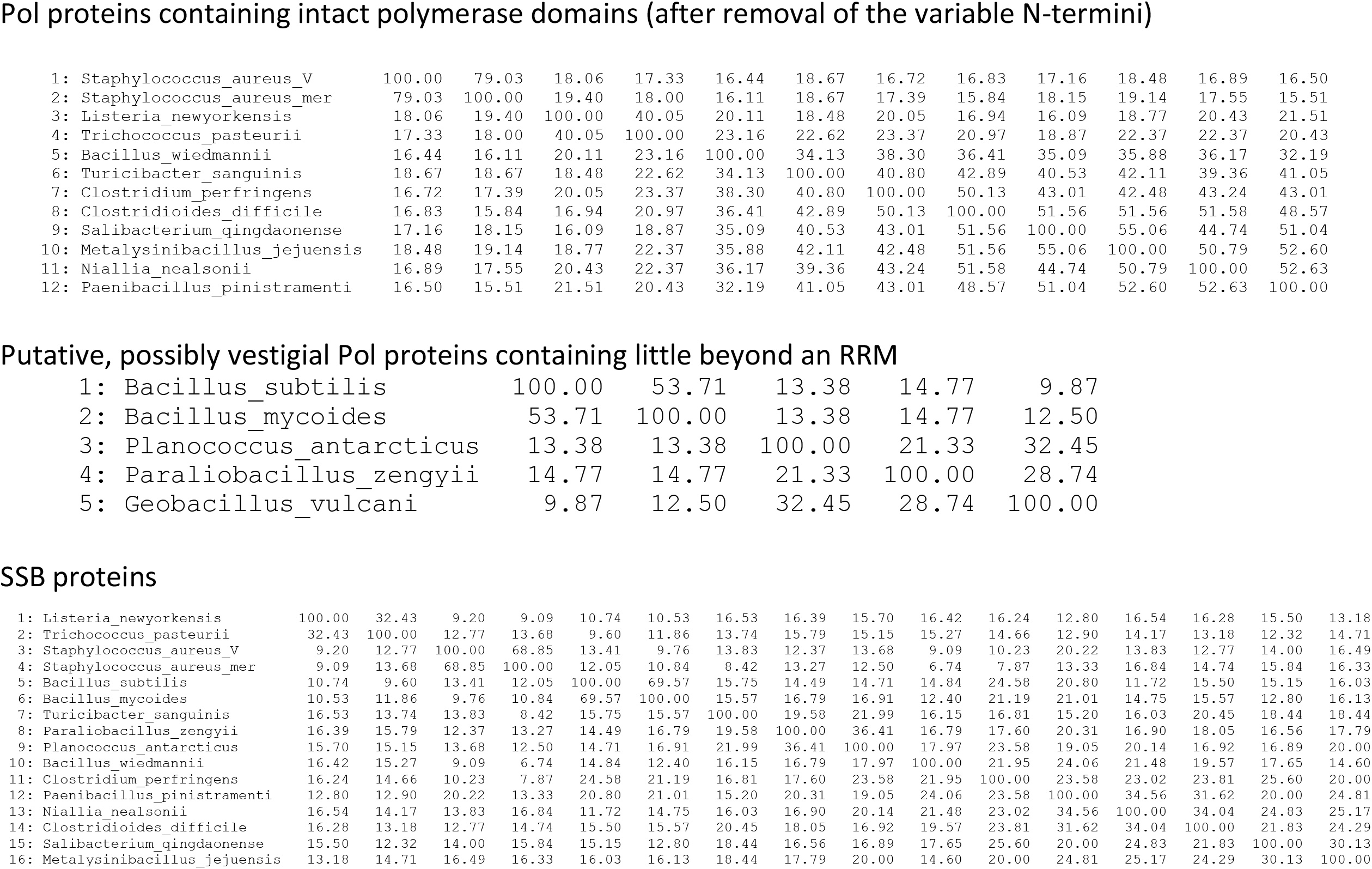
Pairwise sequence identities among proteins studied, as calculated by Clustal Omega

Predicted Alignment Error plots for all predicted complex structures.

In all cases, protein A is the putative polymerase and B is the putative SSB.

Plots are shown for all 5 models predicted for each complex.

*Please see the last page for a guide to interpreting these plots*.

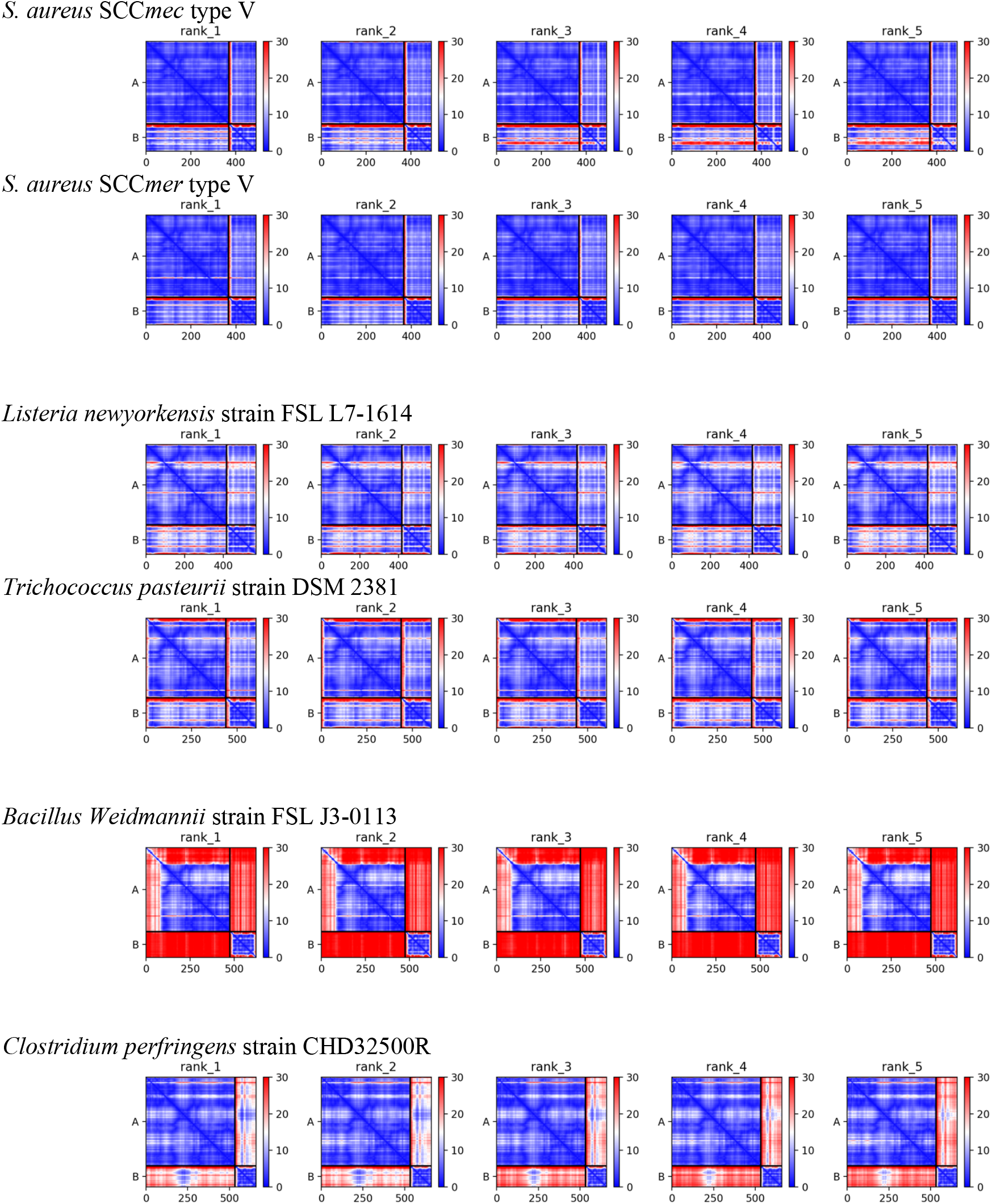
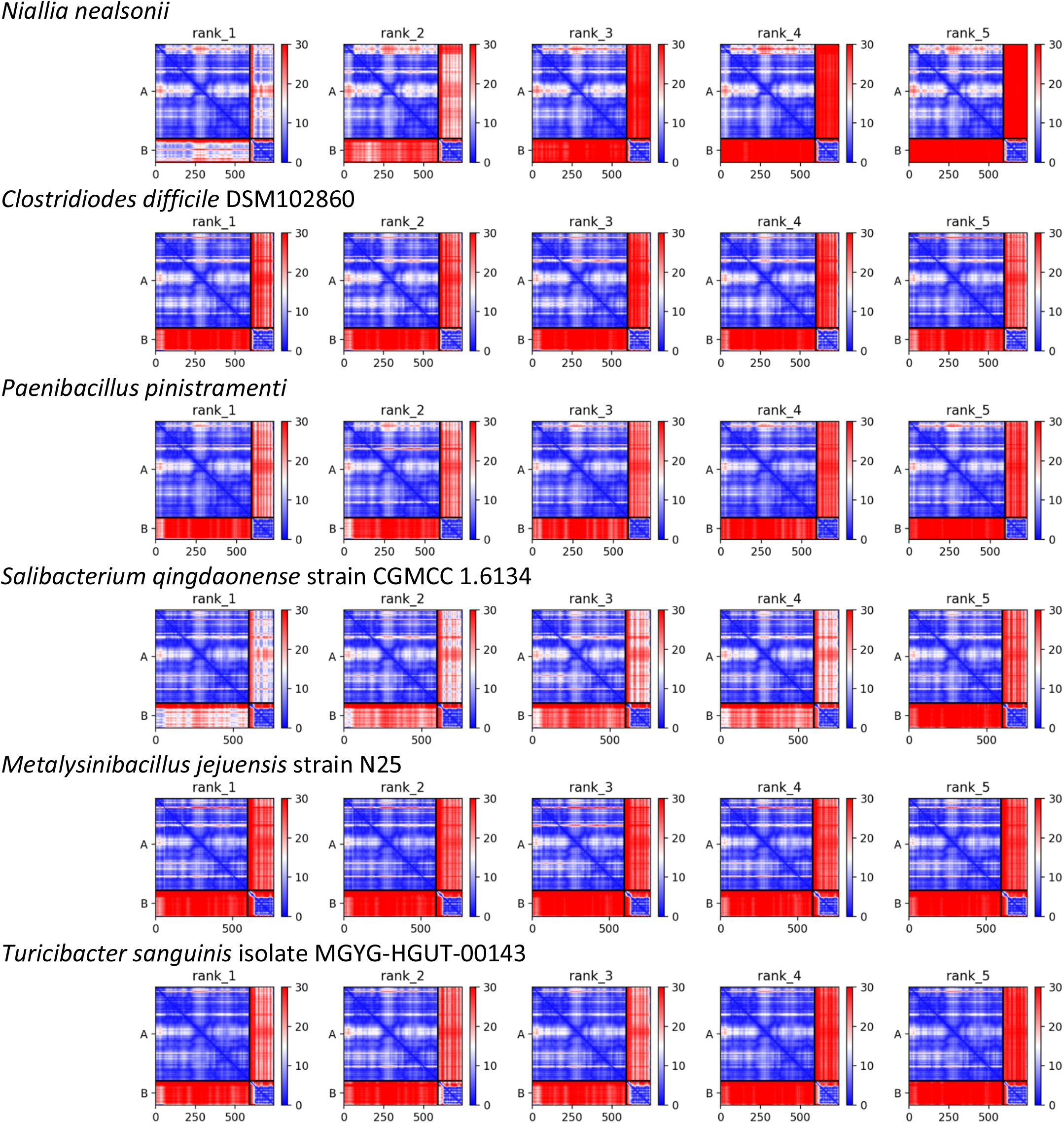

**Putative Pol proteins with little beyond the RRM motif (no protein-protein contacts predicted for any of the following)**

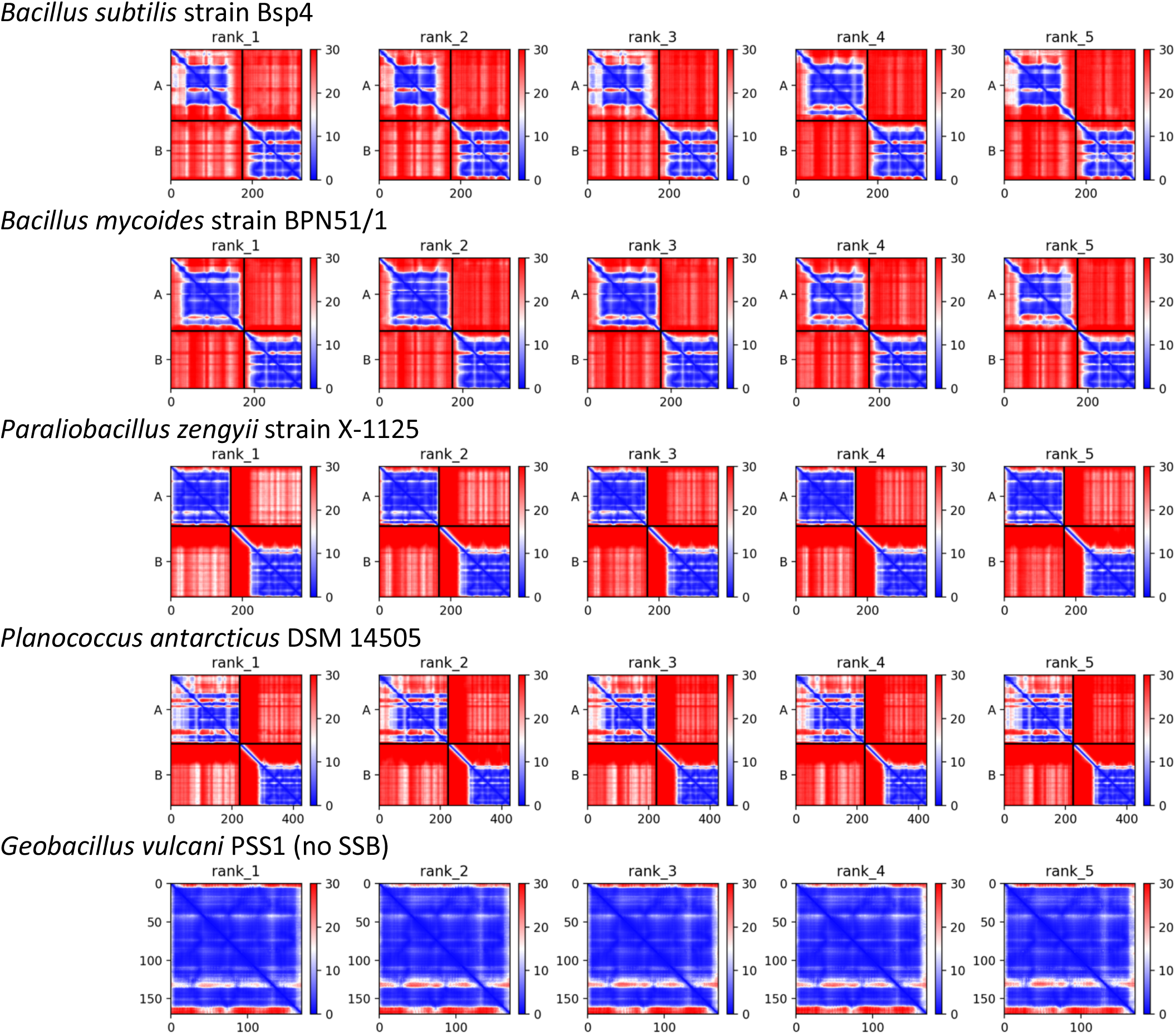

**Guide to interpreting PAE (predicted alignment error plots) from AlphaFold**, using the top model for the *C. difficile* complex as an example. The predicted error in relative positioning between all pairs of residues is color-coded from blue (low) to red (high). On-diagonal boxes reflect the folding of individual proteins (or domains) and off-diagonal boxes reflect the positioning of one protein (or domain) relative to the others. Protein A is the predicted polymerase and B is the predicted SSB. Green boxes highlight the various domains or subdomains of the polymerase. Yellow off-diagonal boxes highlight less-blue sections of the plot which indicate that the coiled coil portion of the thumb is predicted to be connected to the rest of the polymerase by a flexible hinge.

The small off-diagonal green boxes highlight the predicted interaction between the C-terminal extension of the SSB with the N-terminal extension of the polymerase.

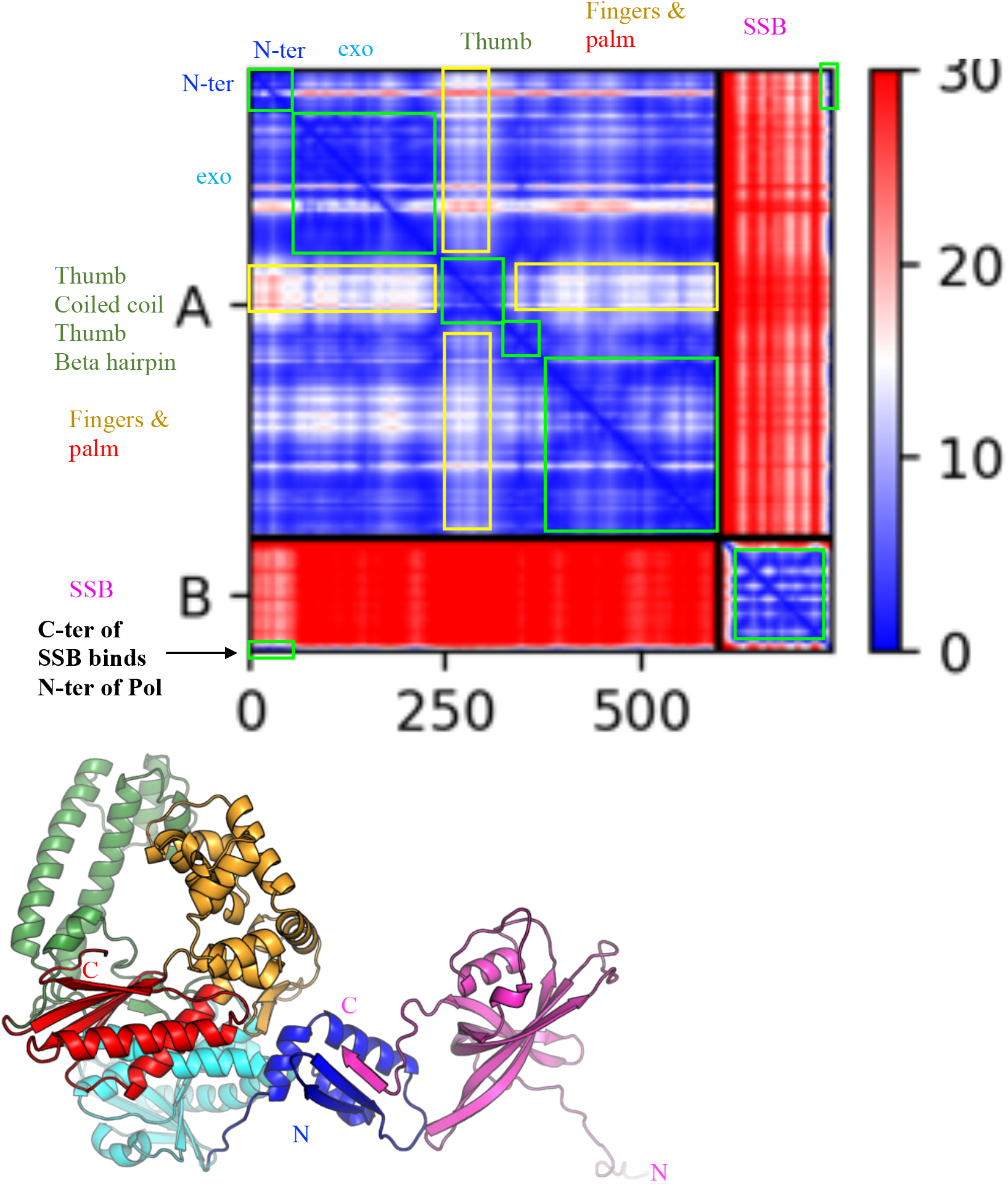

Alignment of all “large” Pol proteins with the 1qsl.pdb - a structure of the Klenow fragment of *E. coli* DNA Pol I, which lacks the 5’-3’ exonuclase but includes the 3’-5’ exonuclease and the polymerase domains.

Promals3D was used to create this alignment.

Active site residues are marked with asterisks over the sequences (blue for the 3’-5’ exonuclease and red for the polymerase sites), and Helix O of the *E. coli* Pol I, which makes important interactions with substrates, is also labeled. The helix that is shorter in the predicted structures, relevant to binding of the 1^st^ dNTP, is marked with #.

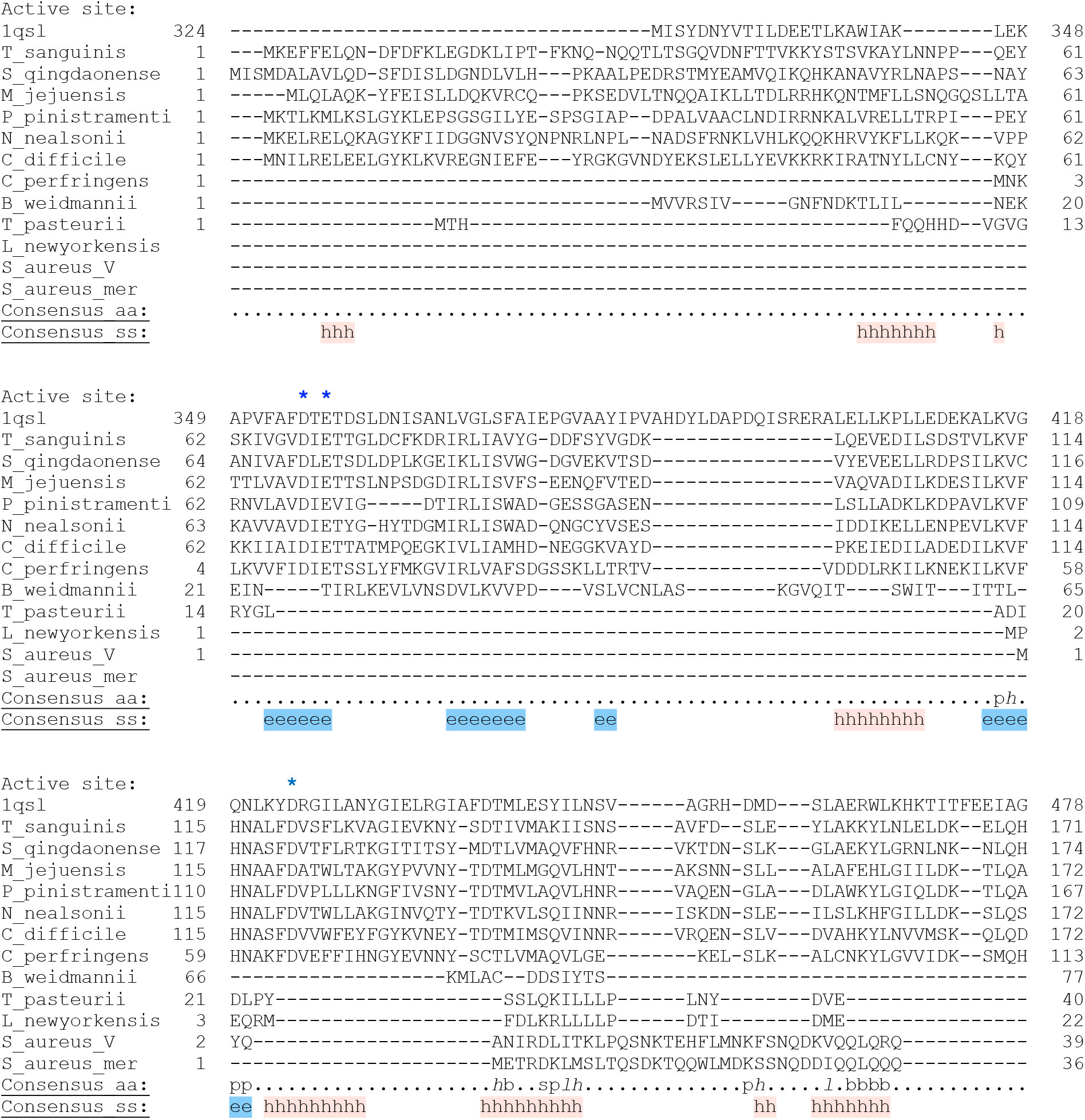
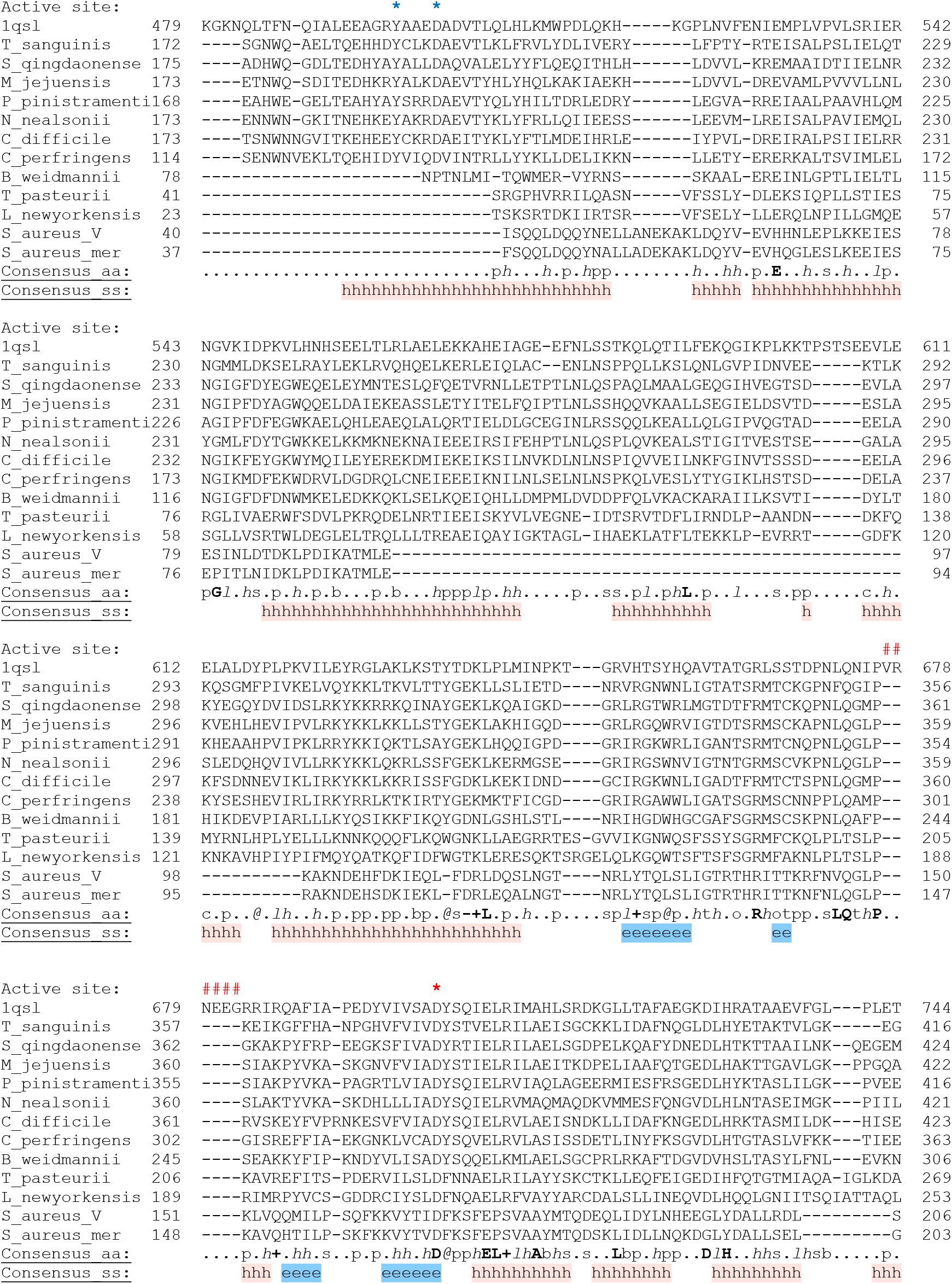
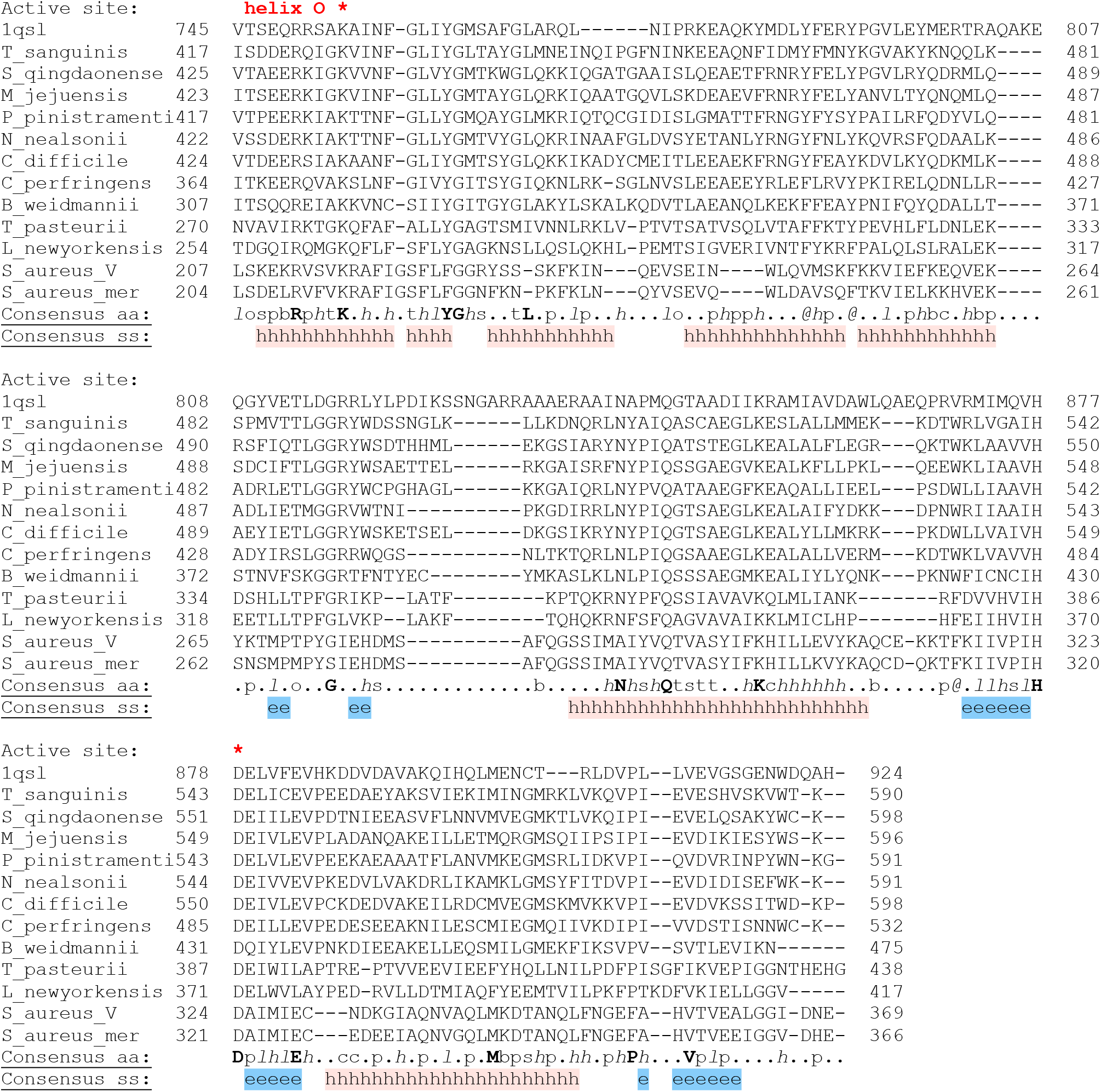

## References

Aravind, L., Leipe, D.D., Koonin, E.V., 1998. Toprim--a conserved catalytic domain in type IA and II topoisomerases, DnaG-type primases, OLD family nucleases and RecR proteins. Nucleic Acids Res. 26, 4205–4213. https://doi.org/10.1093/nar/26.18.4205

Bebel, A., Walsh, M.A., Mir-Sanchis, I., Rice, P.A., 2020. A novel DNA primase-helicase pair encoded by SCCmec elements. eLife 9. https://doi.org/10.7554/eLife.55478

Bedford, E., Tabor, S., Richardson, C.C., 1997. The thioredoxin binding domain of bacteriophage T7 DNA polymerase confers processivity on Escherichia coli DNA polymerase I. Proc. Natl. Acad. Sci. U. S. A. 94, 479–484. https://doi.org/10.1073/pnas.94.2.479

Bergsch, J., Allain, F.H.-T., Lipps, G., 2019. Recent advances in understanding bacterial and archaeoeukaryotic primases. Curr. Opin. Struct. Biol. 59, 159–167. https://doi.org/10.1016/j.sbi.2019.08.004

Blokesch, M., 2016. Natural competence for transformation. Curr. Biol. CB 26, 3255. https://doi.org/10.1016/j.cub.2016.11.023

Brautigam, C.A., Aschheim, K., Steitz, T.A., 1999. Structural elucidation of the binding and inhibitory properties of lanthanide (III) ions at the 3’-5’ exonucleolytic active site of the Klenow fragment. Chem. Biol. 6, 901–908. https://doi.org/10.1016/s1074-5521(00)80009-5

Castro, C., Smidansky, E.D., Arnold, J.J., Maksimchuk, K.R., Moustafa, I., Uchida, A., Götte, M., Konigsberg, W., Cameron, C.E., 2009. Nucleic acid polymerases use a general acid for nucleotidyl transfer. Nat. Struct. Mol. Biol. 16, 212–218. https://doi.org/10.1038/nsmb.1540

Dai, X., Rothman-Denes, L.B., 1998. Sequence and DNA structural determinants of N4 virion RNA polymerase-promoter recognition. Genes Dev. 12, 2782–2790. https://doi.org/10.1101/gad.12.17.2782

Dickey, T.H., Altschuler, S.E., Wuttke, D.S., 2013. Single-stranded DNA-binding proteins: multiple domains for multiple functions. Struct. Lond. Engl. 1993 21, 1074–1084. https://doi.org/10.1016/j.str.2013.05.013

Doublié, S., Tabor, S., Long, A.M., Richardson, C.C., Ellenberger, T., 1998. Crystal structure of a bacteriophage T7 DNA replication complex at 2.2 A resolution. Nature 391, 251–258. https://doi.org/10.1038/34593

Evans, R., O’Neill, M., Pritzel, A., Antropova, N., Senior, A., Green, T., Žídek, A., Bates, R., Blackwell, S., Yim, J., Ronneberger, O., Bodenstein, S., Zielinski, M., Bridgland, A., Potapenko, A., Cowie, A., Tunyasuvunakool, K., Jain, R., Clancy, E., Kohli, P., Jumper, J., Hassabis, D., 2022. Protein complex prediction with AlphaFold-Multimer. https://doi.org/10.1101/2021.10.04.463034

Fillol-Salom, A., Martínez-Rubio, R., Abdulrahman, R.F., Chen, J., Davies, R., Penadés, J.R., 2018. Phage-inducible chromosomal islands are ubiquitous within the bacterial universe. ISME J. 12, 2114–2128. https://doi.org/10.1038/s41396-018-0156-3

Firth, N., Jensen, S.O., Kwong, S.M., Skurray, R.A., Ramsay, J.P., 2018. Staphylococcal Plasmids, Transposable and Integrative Elements. Microbiol. Spectr. 6. https://doi.org/10.1128/microbiolspec.GPP3-0030-2018

Iyer, L.M., Abhiman, S., Aravind, L., 2008. A new family of polymerases related to superfamily A DNA polymerases and T7-like DNA-dependent RNA polymerases. Biol. Direct 3, 39. https://doi.org/10.1186/1745-6150-3-39

Iyer, L.M., Koonin, E.V., Leipe, D.D., Aravind, L., 2005. Origin and evolution of the archaeo-eukaryotic primase superfamily and related palm-domain proteins: structural insights and new members. Nucleic Acids Res. 33, 3875–3896. https://doi.org/10.1093/nar/gki702

Johnson, S.J., Taylor, J.S., Beese, L.S., 2003. Processive DNA synthesis observed in a polymerase crystal suggests a mechanism for the prevention of frameshift mutations. Proc. Natl. Acad. Sci. U. S. A. 100, 3895–3900. https://doi.org/10.1073/pnas.0630532100

Jumper, J., Evans, R., Pritzel, A., Green, T., Figurnov, M., Ronneberger, O., Tunyasuvunakool, K., Bates, R., Žídek, A., Potapenko, A., Bridgland, A., Meyer, C., Kohl, S.A.A., Ballard, A.J., Cowie, A., Romera-Paredes, B., Nikolov, S., Jain, R., Adler, J., Back, T., Petersen, S., Reiman, D., Clancy, E., Zielinski, M., Steinegger, M., Pacholska, M., Berghammer, T., Bodenstein, S., Silver, D., Vinyals, O., Senior, A.W., Kavukcuoglu, K., Kohli, P., Hassabis, D., 2021. Highly accurate protein structure prediction with AlphaFold. Nature. https://doi.org/10.1038/s41586-021-03819-2

Kennedy, W.P., Momand, J.R., Yin, Y.W., 2007. Mechanism for de novo RNA synthesis and initiating nucleotide specificity by t7 RNA polymerase. J. Mol. Biol. 370, 256–268. https://doi.org/10.1016/j.jmb.2007.03.041

Koonin, E.V., Krupovic, M., Ishino, S., Ishino, Y., 2020. The replication machinery of LUCA: common origin of DNA replication and transcription. BMC Biol. 18, 61. https://doi.org/10.1186/s12915-020-00800-9

Maree, M., Thi Nguyen, L.T., Ohniwa, R.L., Higashide, M., Msadek, T., Morikawa, K., 2022. Natural transformation allows transfer of SCCmec-mediated methicillin resistance in Staphylococcus aureus biofilms. Nat. Commun. 13, 2477. https://doi.org/10.1038/s41467-022-29877-2

Minnick, D.T., Astatke, M., Joyce, C.M., Kunkel, T.A., 1996. A thumb subdomain mutant of the large fragment of Escherichia coli DNA polymerase I with reduced DNA binding affinity, processivity, and frameshift fidelity. J. Biol. Chem. 271, 24954–24961. https://doi.org/10.1074/jbc.271.40.24954

Mirdita, M., Schütze, K., Moriwaki, Y., Heo, L., Ovchinnikov, S., Steinegger, M., 2022. ColabFold: making protein folding accessible to all. Nat. Methods 19, 679–682. https://doi.org/10.1038/s41592-022-01488-1

Mir-Sanchis, I., Roman, C.A., Misiura, A., Pigli, Y.Z., Boyle-Vavra, S., Rice, P.A., 2016. Staphylococcal SCCmec elements encode an active MCM-like helicase and thus may be replicative. Nat. Struct. Mol. Biol. 23, 891–898. https://doi.org/10.1038/nsmb.3286

Misiura, A., Pigli, Y.Z., Boyle-Vavra, S., Daum, R.S., Boocock, M.R., Rice, P.A., 2013. Roles of two large serine recombinases in mobilizing the methicillin-resistance cassette SCCmec. Mol. Microbiol. 88, 1218–1229. https://doi.org/10.1111/mmi.12253

Pei, J., Kim, B.-H., Grishin, N.V., 2008. PROMALS3D: a tool for multiple protein sequence and structure alignments. Nucleic Acids Res. 36, 2295–2300. https://doi.org/10.1093/nar/gkn072

Polesky, A.H., Dahlberg, M.E., Benkovic, S.J., Grindley, N.D., Joyce, C.M., 1992. Side chains involved in catalysis of the polymerase reaction of DNA polymerase I from Escherichia coli. J. Biol. Chem. 267, 8417–8428.

Qiao, C., Debiasi-Anders, G., Mir-Sanchis, I., 2022. Staphylococcal self-loading helicases couple the staircase mechanism with inter domain high flexibility. Nucleic Acids Res. 50, 8349–8362. https://doi.org/10.1093/nar/gkac625

Raghunathan, S., Kozlov, A.G., Lohman, T.M., Waksman, G., 2000. Structure of the DNA binding domain of E. coli SSB bound to ssDNA. Nat. Struct. Biol. 7, 648–652. https://doi.org/10.1038/77943

Raia, P., Delarue, M., Sauguet, L., 2019. An updated structural classification of replicative DNA polymerases. Biochem. Soc. Trans. 47, 239–249. https://doi.org/10.1042/BST20180579

Rocha, E.P.C., Bikard, D., 2022. Microbial defenses against mobile genetic elements and viruses: Who defends whom from what? PLoS Biol. 20, e3001514. https://doi.org/10.1371/journal.pbio.3001514

Saha, C.K., Sanches Pires, R., Brolin, H., Delannoy, M., Atkinson, G.C., 2021. FlaGs and webFlaGs: discovering novel biology through the analysis of gene neighbourhood conservation. Bioinforma. Oxf. Engl. 37, 1312–1314. https://doi.org/10.1093/bioinformatics/btaa788

Shen, Z., Tang, C.M., Liu, G.-Y., 2022. Towards a better understanding of antimicrobial resistance dissemination: what can be learnt from studying model conjugative plasmids? Mil. Med. Res. 9, 3. https://doi.org/10.1186/s40779-021-00362-z

Shereda, R.D., Kozlov, A.G., Lohman, T.M., Cox, M.M., Keck, J.L., 2008. SSB as an organizer/mobilizer of genome maintenance complexes. Crit. Rev. Biochem. Mol. Biol. 43, 289–318. https://doi.org/10.1080/10409230802341296

Sievers, F., Wilm, A., Dineen, D., Gibson, T.J., Karplus, K., Li, W., Lopez, R., McWilliam, H., Remmert, M., Söding, J., Thompson, J.D., Higgins, D.G., 2011. Fast, scalable generation of high-quality protein multiple sequence alignments using Clustal Omega. Mol. Syst. Biol. 7, 539. https://doi.org/10.1038/msb.2011.75

Ubeda, C., Barry, P., Penadés, J.R., Novick, R.P., 2007. A pathogenicity island replicon in Staphylococcus aureus replicates as an unstable plasmid. Proc. Natl. Acad. Sci. U. S. A. 104, 14182–14188. https://doi.org/10.1073/pnas.0705994104

## References

PROMALS3D: a tool for multiple sequence and structure alignment. Jimin Pei, Bong-Hyun Kim and Nick V. Grishin. Nucleic Acids Res. 2008 36(7):2295–2300.

PDB ID 1qsl: Structural elucidation of the binding and inhibitory properties of lanthanide (III) ions at the 3’-5’ exonucleolytic active site of the Klenow fragment.

Brautigam CA, Aschheim K, Steitz TA. Chem Biol. 1999 Dec;6(12):901–8. doi: 10.1016/s1074-5521(00)80009-5. PMID: 10631518

